# Modeling Cystic Fibrosis Chronic Infection Using Engineered Mucus-like Hydrogels

**DOI:** 10.1101/2023.09.07.556675

**Authors:** Courtney L. O’Brien, Sarah Spencer, Naeimeh Jafari, Andy J. Huang, Alison J. Scott, Zhenyu Cheng, Brendan M. Leung

## Abstract

The airway mucus of patients with cystic fibrosis has altered properties which create a microenvironment primed for chronic infections that are difficult to treat. These complex polymicrobial airway infections and corresponding mammalian-microbe interactions are challenging to model *in vitro*. Here, we report the development of mucus-like hydrogels with varied compositions and viscoelastic properties reflecting differences between healthy and cystic fibrosis airway mucus. Models of cystic fibrosis and healthy airway microenvironments were created by combining the hydrogels with relevant pathogens, human bronchial epithelial cells, and an antibiotic. Notably, pathogen antibiotic resistance was not solely dependent on the altered properties of the mucus-like hydrogels but was also influenced by culture conditions including microbe species, monomicrobial or polymicrobial culture, and the presence of epithelial cells. Additionally, the cystic fibrosis airway model showed the ability to mimic features characteristic of chronic cystic fibrosis airway infections including sustained polymicrobial growth and increased antibiotic tolerance.

## Introduction

Airway mucus acts as a protective barrier for the respiratory system by collecting and clearing harmful pathogens from the airways^1,2^. In cystic fibrosis (CF), mutations of the *cystic fibrosis transmembrane conductance regulator* (*CFTR*) gene cause the airway epithelial surface to be dehydrated, leading to increased solids and mucin protein concentrations and altered mucus viscoelasticity^3–7^. These altered airway mucus properties impair mucus clearance mechanisms such as mucociliary and cough mediated clearance ^8–10^. Impaired mucus clearance in CF results in mucus layer thickening and chronic polymicrobial airway infections as opportunistic pathogens, such as *Pseudomonas aeruginosa* and *Staphylococcus aureus*, colonize the stagnant airway mucus. Pathogens that infect the airways of CF patients are also prone to developing resistance to antibiotics which makes these infections increasingly difficult to treat^11,12^. Respiratory function decline due to chronic airway infections is the leading cause of mortality in CF patients^13^. This emphasizes the importance of investigating microbe-microbe and host-microbe interactions within the epithelial mucosal niche.

*In vivo* CF models, which include small animal models, such as mice^14,15^, and larger animal models, such as pigs^16,17^, are limited in their ability to be used to study CF chronic infections due to physiological differences and extensive resource requirements. Due to these limitations, many *in vitro* CF airway models have been developed and explored. A number of these *in vitro* studies have investigated interactions between mammalian cells and *P. aeruginosa* and/or *S. aureus*^18–21^, but without a mucus-like layer in these systems, they are not able to fully represent interactions within the CF airway mucosal niche. Other *in vitro* CF models focus on the growth behavior of CF pathogens when cultured with CF mucus or mucus-mimetics without the inclusion of mammalian cells^22–25^. Mammalian cells are an important inclusion within an *in vitro* CF airway model as they allow for the observation of two-way host-microbe interactions and render the model more representative of the native CF airway microenvironment.

Artificial CF sputum medium (ASM)^22,26^ and synthetic CF sputum medium (SCFM)^27^ are two commonly used synthetic mucus mimetic materials for model systems. Although their components provide the chemical constituents relevant to the CF airway, neither of these materials have the hallmark viscoelastic properties of CF airway mucus. Both ASM and modified SCFM have shown viscous behavior rather than the viscoelastic solid behavior that is characteristic of CF airway mucus^28,29^. The combination of porcine gastric mucins (PGM) with gel-forming polymers is another method used to control the viscoelasticity of mucus model systems^30^. For example, this can be achieved by combining PGM with natural polymers, such as hyaluronic acid^31^ or sodium alginate^32–34^. Additionally, many of the models that do include mucus or a mucus-like component only model monomicrobial infections rather than modeling the more complex polymicrobial interactions that are known to occur within the CF airways.

The limitations of current CF models creates the need for an *in vitro* polymicrobial infection model of the CF microenvironment with a simple yet representative mucus-like component. In this study, two mucus-like hydrogels were developed to capture key relative differences between CF and healthy airway mucus, including their distinct solids concentration and altered viscoelastic properties. These hydrogels include a crosslinked gel component to obtain mucus-like viscoelastic properties and a mucin protein component to capture one of the major changes in composition between healthy and CF airway mucus. These hydrogels were overlaid on human bronchial epithelial cells (16HBE14o-) to recapitulate healthy and CF airways. The airway models consist of relevant CF pathogens, *P. aeruginosa* and *S. aureus*, were cultured over top of the mucus-like hydrogels in the presence of ciprofloxacin, a relevant CF antibiotic (Figure 1).

**Figure 1:**
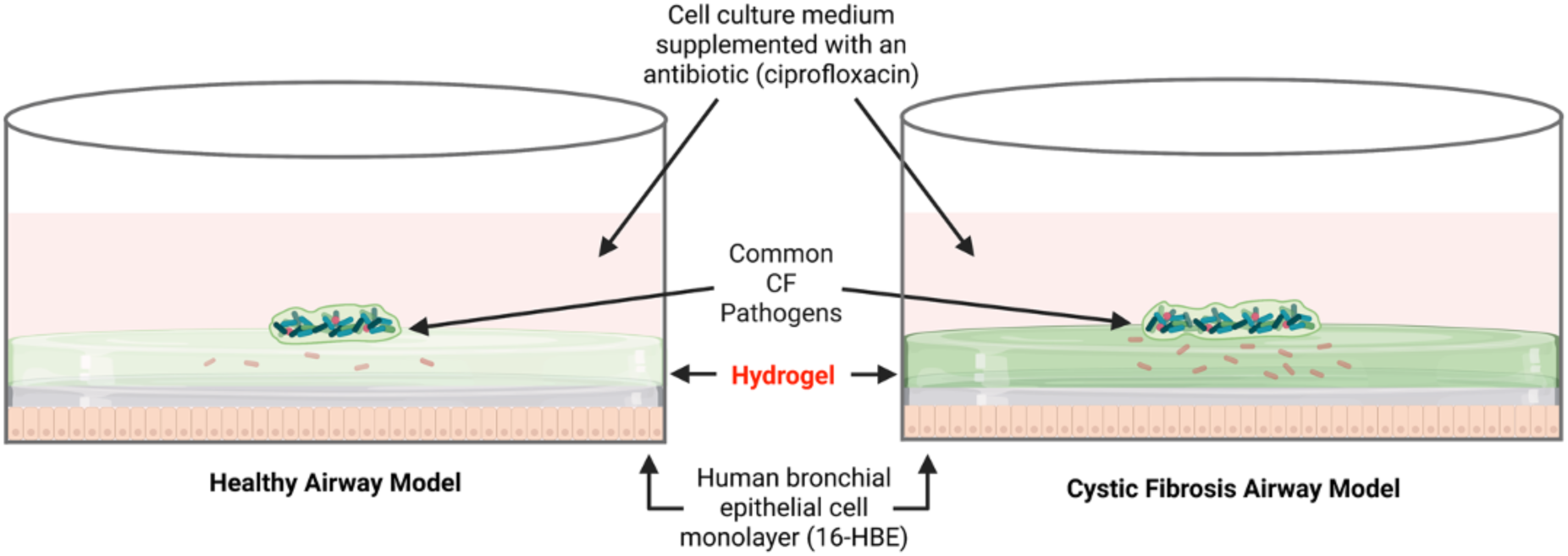
Schematic of the healthy and CF airway model. The model includes cells from a human bronchial epithelial cell line (16HBE), mucus-like hydrogels, common CF pathogens, and cell culture medium supplemented with a relevant CF antibiotic (ciprofloxacin). The healthy airway model (left) includes the healthy mucus-like hydrogel and the CF airway model (right) includes the CF mucus-like hydrogel.

We showed that the altered compositions of these mucus-like hydrogels can influence hydrogel viscoelasticity, mammalian cell viability, bacterial growth behavior, and antibiotic tolerance. In addition, the CF mucus-like hydrogel enabled us to extend the longevity of the model system to surpass current culture platforms and better capture aspects of chronic CF infections^20,35–38^. The healthy and CF *in vitro* airway models developed in this research are useful for observing mono- and polymicrobial behavior and response to relative changes within their microenvironment. This can provide a better understanding of how the physical properties and composition of airway mucus can impact cell behaviors within the microbe-mucosal-mammalian niche.

## Results

### Alginate-based mucus-like hydrogels mimic the viscoelastic properties of physiological airway mucus and support mammalian cell growth

To generate *in vitro* models of the healthy and CF airway microenvironments, we developed a number of potential mucus-like hydrogels using alginate as the gel forming component with the addition of mucins to confer physiological functions (Table S1). Compositions including various concentrations of alginate, crosslinker (calcium chloride), and mucin were developed to represent the altered viscoelasticity and solid concentrations of the CF mucus. The viscoelastic properties of the mucus-like hydrogels were characterized through measurement of their storage moduli and loss moduli (Table S1) across a range of angular frequencies between 0.1 and 20 rad/s (Figure 2). The viscoelastic measurements of the hydrogels were compared to measured values of collected mucus samples reported in literature, two gels were chosen to be suitable candidates to represent CF and healthy mucus. The healthy mucus-like hydrogel formulation consisted of 1.5% w/v alginate, 0.5 mg/ml CaCl_2_, and 1% w/v mucins. The CF mucus-like hydrogel formulation consisted of 1.5% alginate, 0.6 mg/ml CaCl_2_, and 4% mucins. In comparing the two mucus-like hydrogel responses at each measured angular frequency, the average storage and loss moduli were at least 1.6 times higher for the CF mucus-like hydrogel, relative to the healthy mucus-like hydrogel counterpart.

**Figure 2:**
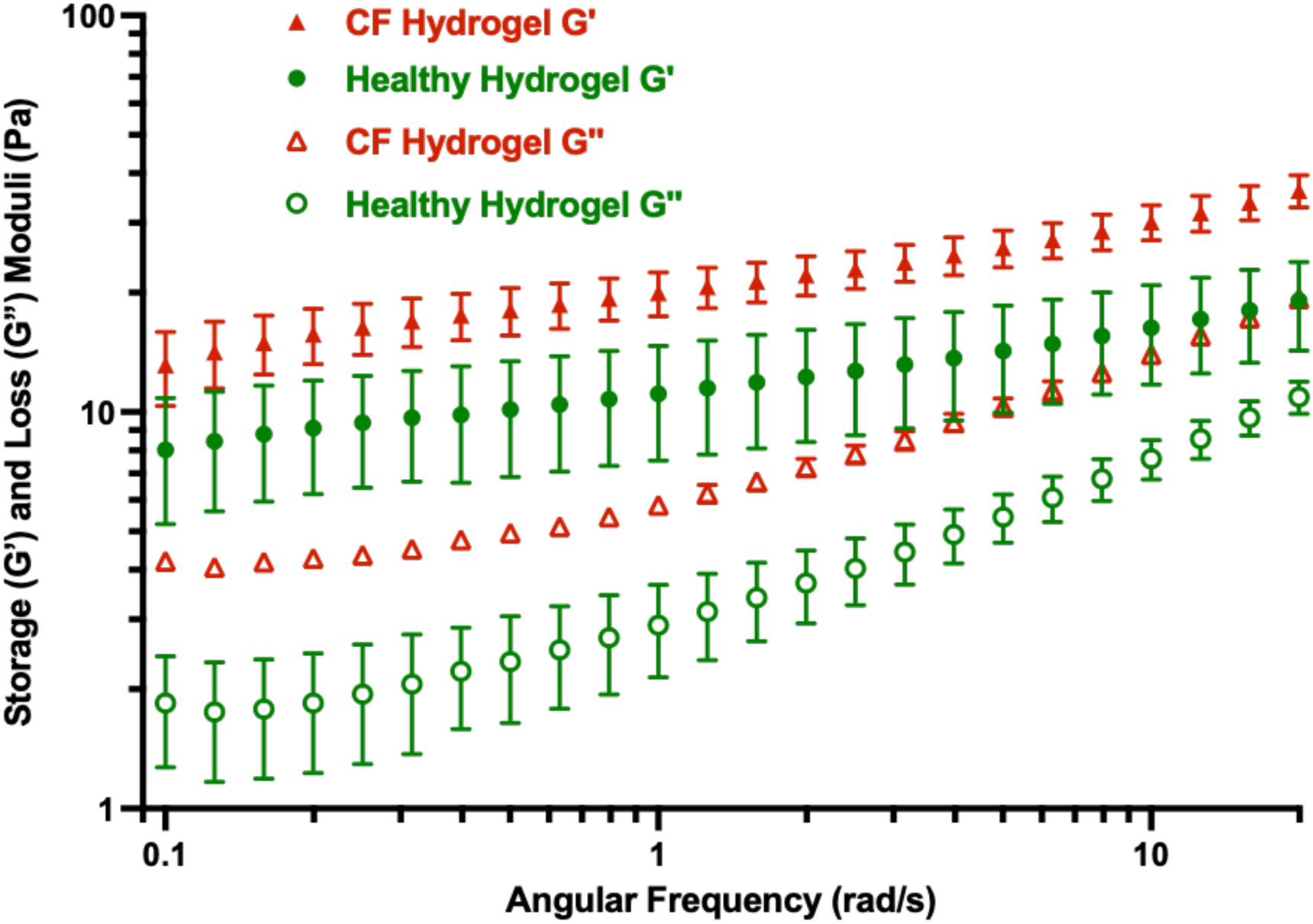
*Viscoelastic properties of the chosen mucus-like hydrogel formulations.* The properties of the CF mucus-like hydrogel are shown in red and the properties of the healthy mucus-like hydrogel are shown in green. Storage moduli, G’, represented by solid icons and loss moduli, G’’, represented by open icons. Data shown from an angular frequency range of 0.1 to 20 rad/s. Values are the average ± SD from three independent measurements.

The viability of 16HBE cells was assessed after 48 hours of culture with the CF and healthy alginate hydrogels. A total hydrogel thickness of 1.5 mm was chosen to approximate a build-up of mucus in the bronchi/bronchioles, which has been found to range from 1 to approximately 6 mm^39^. When 16HBE cells were cultured in the presence of the mucus-like hydrogels, cell viability was found to be 98% or greater for both the CF and healthy mucus-like hydrogel culture conditions (Figure 3).

**Figure 3:**
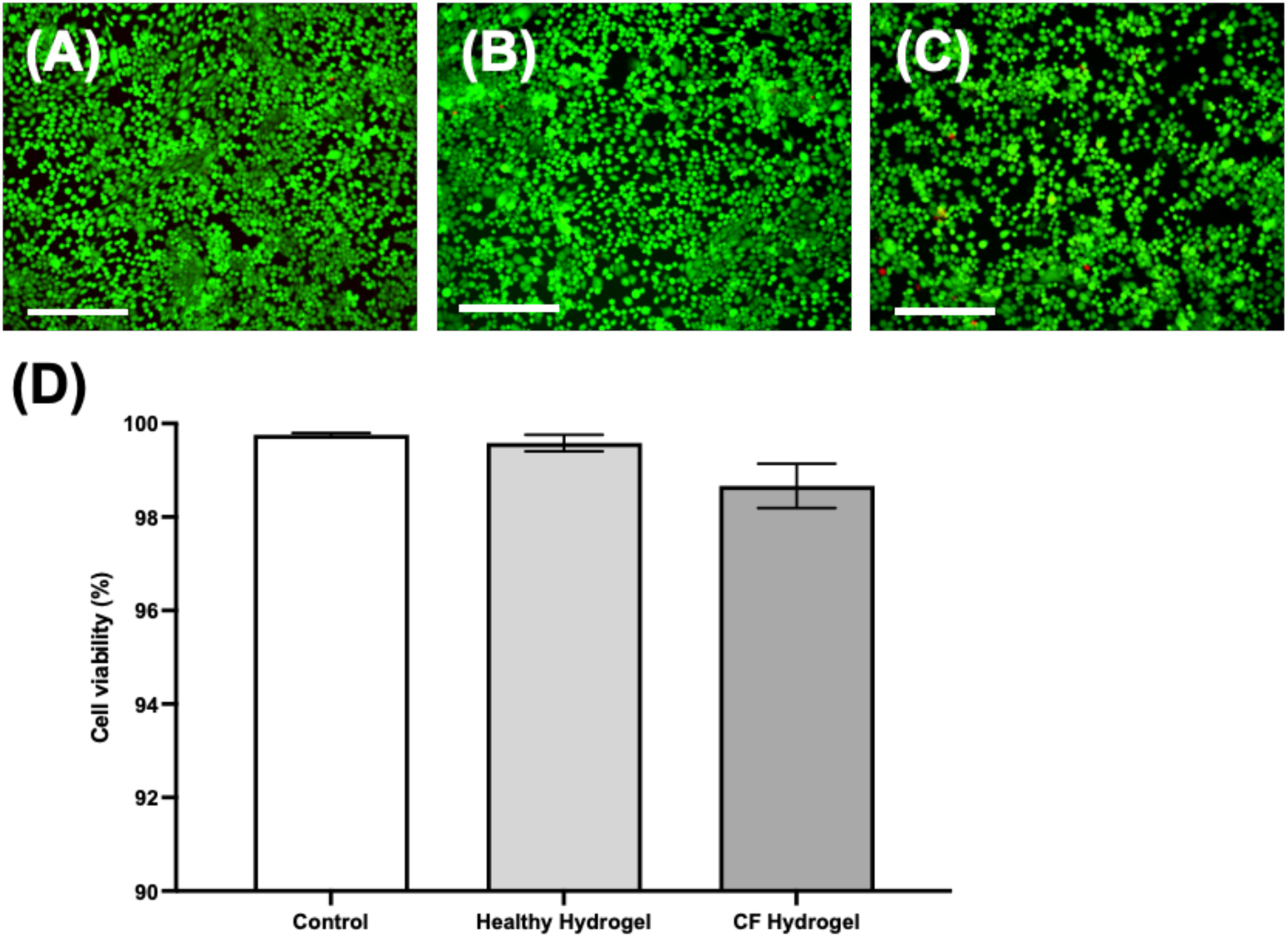
Cell viability of 16HBE cells cultured beneath CaCl_2_-crosslinked alginate hydrogels with the addition of an alginate cushion layer. Live/dead assay performed after 48 hours of culture. Green cells are viable and red cells are non-viable. **(A)** Control, 16HBE cells grown with no hydrogel. **(B)** 16HBE cells grown with the healthy mucus-like hydrogel. **(C)** 16HBE cells grown with the CF mucus-like hydrogel. Representative images chosen from three biological replicates. Scale bar is 275 μm. **(D)** Average 16HBE cell viability (%) ± SD. Data averaged from three biological replicates.

### Bacteria growth and antibiotic susceptibility is impacted by the properties of mucus-like hydrogels within *in vitro* polymicrobial airway models

The minimum inhibitory concentration (MIC) of ciprofloxacin against *P. aeruginosa* and *S. aureus* was determined to be 0.125 μg/ml. A ciprofloxacin concentration of 4x MIC (0.5 μg/ml) was used for this study. An aqueous two-phase system (ATPS) was used to confine and establish patterned growth of *P. aeruginosa* and *S. aureus* in monomicrobial and polymicrobial form on top of the mucus-like hydrogels. After 5 hours, minimal viable bacteria were detected within the liquid phase of the culture (i.e., PEG phase of the ATPS) (Figure S1A-D). In monoculture, both *P. aeruginosa* and *S. aureus* showed increased resistance to ciprofloxacin when grown with the CF mucus-like hydrogel compared to the healthy mucus-like hydrogel (Figure 4A and 4C). When cultured with *S. aureus*, *P. aeruginosa* also showed increased resistance to ciprofloxacin when grown with the CF hydrogel (Figure 4E), this trend was observed for all replicates. Interestingly, the antibiotic resistance of *S. aureus* conferred when cultured with the CF mucus hydrogel was lost when cultured together with *P. aeruginosa*.

**Figure 4:**
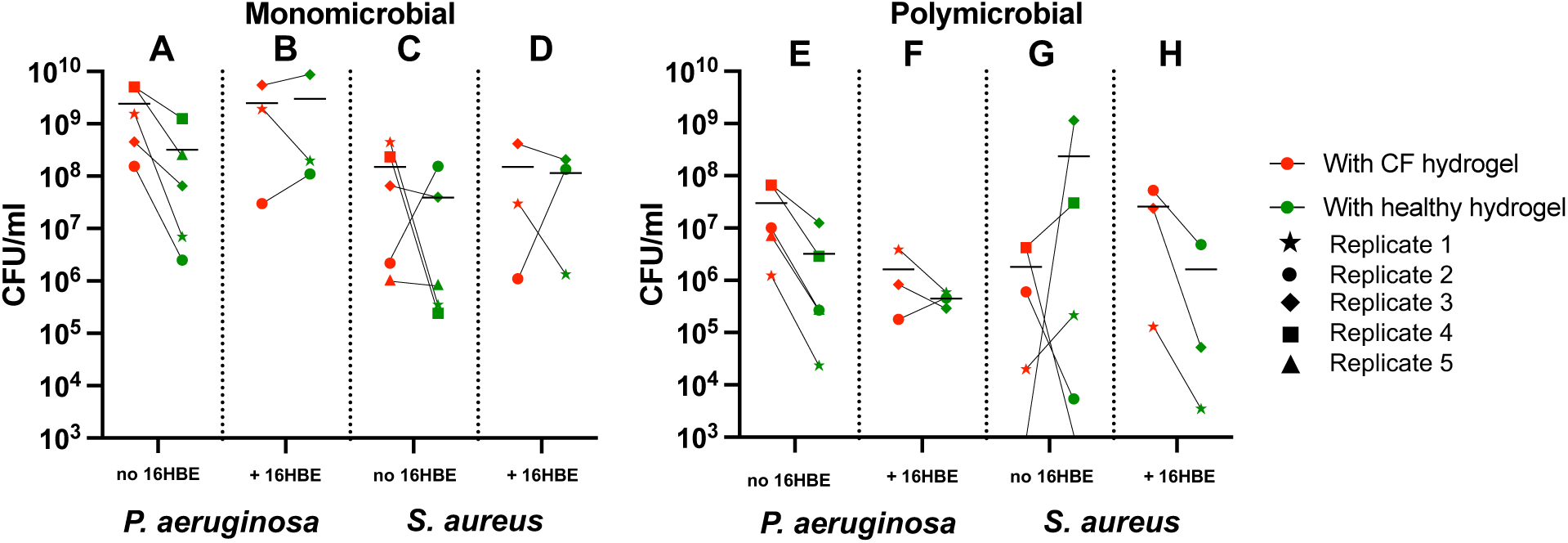
Concentrations of viable bacteria grown with healthy and CF mucus-like hydrogels after 48 hours in the presence of 0.5 μg/ml ciprofloxacin. **(A)** P. aeruginosa and *S. aureus* grown in monomicrobial conditions without and with 16HBE cells and **(B)** P. aeruginosa and *S. aureus* grown in polymicrobial conditions without and with 16HBE cells. A different icon indicates the values from each biological replicate, and horizontal black bars represent the mean (n = 5 for experiments without 16HBE cells and n = 3 for experiments with 16HBE cells). Values of growth with the CF and healthy hydrogels from the same biological replicate are connected by a line. Any missing icons or lines passing through the x-axis indicate a value of zero.

The inclusion of 16HBE cells led to observable changes in the pattern of antibiotic resistance within the airway models. In monoculture of both *P. aeruginosa* and *S. aureus,* the presence of 16HBE increased their antibiotic resistance within the healthy gel to a level comparable to the CF gel, while antibiotic resistance was not altered within the CF gel compared to 16HBE free culture (Figure 4A-D). In coculture, the presence of 16HBE decreased the antimicrobial resistance of *P. aeruginosa* in CF gel (>10-fold decrease in microbial load) and to a lesser degree in the healthy gel (<10-fold decrease in microbial load) (Figure 4E and F). The presence of 16HBE negates the *P. aeruginosa* antimicrobial resistance conferred by the CF gel compared to healthy gel in both mono and polymicrobial (Figure 4A, B, E, and F). In addition, *P. aeruginosa* became less resistant to antibiotic when cultured with *S. aureus* (∼10-fold decrease in viable bacteria load) regardless of CF or healthy gel (Figure 4E and F).

In contrast, the presence of the CF gel did not confer antimicrobial resistance to monomicrobial *S. aureus* culture as compared to the healthy gel, irrespective of the presence of 16HBE. However, when cultured with *P. aeruginosa* the antibiotic resistance of *S. aureus* was dependent on gel type and the presence of 16HBE (Figure 4G and H). In the absence of 16HBE, *S. aureus* in the healthy gel showed increased resistance to antibiotic as compared to the CF gel, indicated by a 10^3^-fold increase in viable microbial load (Figure 4G). In the presence of 16HBE the opposite was observed where *S. aureus* in the CF gel was more resistant to antibiotics than in the healthy gel, although the difference was less drastic compared to 16HBE-free polymicrobial culture (Figure 4G and H). Also, the absence of 16HBE led to an increase in variance of *S. aureus* antibiotic resistance in the polymicrobial models, in particular when cultured with the healthy gel (Figure 4G).

### Bacteria culture conditions impact mammalian cell viability within *in vitro* co-culture airway models

16HBE cell viability after culture within the healthy and CF airway models for 48 hours was varied between bacteria culture conditions. 16HBE cell viability in the monomicrobial *P. aeruginosa* culture conditions was consistently similar to the bacteria-free control (Figure 5A and 5B). 16HBE cell viability was diminished in the monomicrobial *S. aureus* culture conditions in both the CF and healthy airway models (Figure 5C). The viability of 16HBE cells grown in the polymicrobial airway model falls between what was observed for the two species in monomicrobial culture conditions (Figure 5D). The presence of *S. aureus*, both in monomicrobial and polymicrobial culture conditions, appeared to increase variation in 16HBE cell viability within the models (Figure S2). Cell viability was similar between the CF and healthy airway models, although 16HBE cell confluency was generally reduced for the CF airway models relative to the healthy airway models. An additional observation for the 16HBE cells after culture within the airway models was that, for some of the monomicrobial and polymicrobial *S. aureus* culture conditions, 16HBE cells did not stain as viable (i.e., green) or non-viable (i.e., red), but did stain with the DNA stain (i.e., blue). For these conditions where cellular material was remaining but did not stain as live or dead, an image presenting the Hoechst DNA stain was provided in place of the live/dead image (Figure 5).

**Figure 5:**
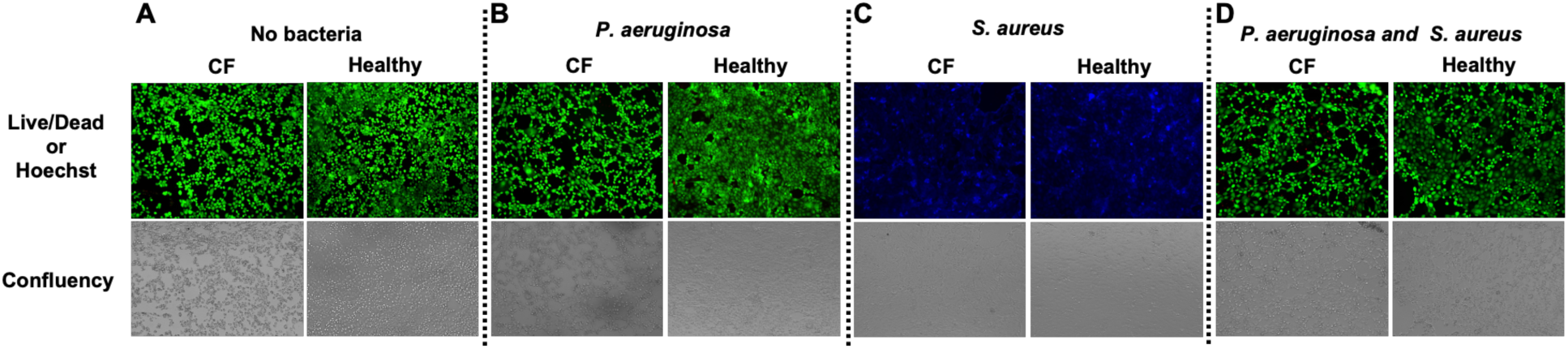
16HBE cell viability and overall adherence after culture within the CF and healthy airway models. Conditions included control models with no bacteria, *P. aeruginosa* monoculture, *S. aureus* monoculture, and *P. aeruginosa* and *S. aureus* co-culture. All conditions were exposed to 0.5 μg/ml ciprofloxacin. Live/dead assay was performed after 48 hours of culture. Green cells are viable and red cells are non-viable. Hoechst DNA stain (blue) presented in conditions where cellular material is remaining but is not staining green or red. Representative images selected from replicate two. Scale bar is 275 μm.

## Discussion

Impaired mucus clearance in CF leads to mucus layer thickening and chronic airway infections as opportunistic pathogens, such as *P. aeruginosa* and *S. aureus*, colonize the stagnant airway mucus. Pathogens that infect the airways of CF patients are also prone to developing resistance to antibiotics which makes these infections increasingly difficult to treat^11,12^. Respiratory function decline due to chronic airway infections is the leading cause of mortality in CF patients^13^ which emphasizes the importance of investigating microbe-microbe and host-microbe interactions in these airway infections. Limitations with *in vivo* animal models of CF airway infections create the need for *in vitro* models of the CF airway microenvironment. Many current *in vitro* models of the CF airway include only one or two of the main elements of this microenvironment, such as microbes/airway cells or microbes/mucus. These models lack the combined components required to capture complex interactions within the CF airway microenvironment. In this study, we developed CF and healthy mucus-like hydrogels which capture the relative differences in viscoelastic properties and component concentrations seen in CF and healthy airway mucus. These mucus-like hydrogels were cultured with human bronchial epithelial cells, relevant CF pathogens, and a common antibiotic used in CF treatment to observe host-microbe and microbe-microbe interactions in response to some of the key differences between the CF and healthy airway microenvironments.

Viscoelastic properties, such as the storage and loss moduli, are obtained through oscillatory testing and are commonly used to characterize healthy and pathologic airway mucus samples in literature. Hydrogel formulations containing 1% and 4% (w/v) mucin proteins were developed to represent healthy and CF mucus-like hydrogels, respectively^4^. These hydrogel formulations are minimalistic by design, while also capable of capturing the relative increase in overall solids concentration as seen in CF mucus compared to healthy airway mucus. Storage and loss moduli of the mucus-like hydrogel formulations were influenced by crosslinker (calcium chloride), alginate, and mucin concentrations. The mucus-like hydrogels showed many features characteristic of physiological airway mucus, most notably, the elastically dominant behavior with storage moduli larger than their respective loss moduli^40–43^. The prevailing trend from studies performed thus far indicates CF airway mucus has greater viscoelastic moduli than healthy airway mucus^6,7^. Additionally, the viscoelastic properties of the CF and healthy mucus-like hydrogels developed herein were within the same order of magnitude as the viscoelastic properties of CF and healthy airway mucus that have been analyzed and reported in literature. Recent sources have found that healthy and CF airway mucus have storage moduli within the range of 0.2 Pa to 93 Pa and loss moduli within the range of 0.1 Pa to 19 Pa^6,7^. The hydrogels developed here recapitulate the key differences in viscoelastic properties, mucin concentration, and solids concentration in physiological CF and healthy airway mucus.

To confirm the non-cytotoxic nature of the healthy and CF mucus-like hydrogels, 16HBE cells were cultured with these gels over 48 hours, and the viabilities were found to be greater than 98% (Figure 3). When *P. aeruginosa* and *S. aureus* were introduced into the healthy and CF airway models without ciprofloxacin, 16HBE viability was assessed after 6 and 24 hours (1 hour and 19 hours, respectively, after removal of PEG-rich phase of the ATPS). The assessment revealed that without the inclusion of antibiotics in the culture system after 6 hours cell viability was generally not impacted but after 24 hours no viable cells remained (data not shown here). As expected, the inclusion of 0.5 μg/ml ciprofloxacin resulted in varied 16HBE cell viability and increased cell survival after 48 hours (Figure 5), suggesting that 16HBE death in subsequent experiments was caused primarily by bacteria and not hydrogel cytotoxicity.

*P. aeruginosa* in culture with *S. aureus* showed increased resistance to ciprofloxacin within the CF airway model as compared to the healthy model (Figure 4F). This trend was also observed between the healthy and CF conditions of *P. aeruginosa* in monoculture without 16HBE cells. The presence of *S. aureus* in co-culture may have enhanced the ability for *P. aeruginosa* to form antibiotic tolerant aggregates or biofilms, as seen in other studies with *P. aeruginosa* cultured with *S. aureus* or *S. aureus* exoproducts^44,45^, especially in mucin-rich microenvironments^23,46^. When 16HBE cells were included within the model in polymicrobial conditions, the concentration of viable *P. aeruginosa* in the healthy and CF hydrogel phases was reduced approximately 10-fold. This observation could be attributed to the presence of antimicrobial peptides (AMPs) produced by epithelial cells, such as human β-defensins (hBDs), the expression of which can be induced by bacteria and bacteria endotoxins^47^. Increased expression of hBDs in 16HBE cells has been observed in the presence of *Aspergillus fumigatus*, a respiratory infection-causing fungus^48^. *S. aureus* has been found to increase expression of epithelial hBD-2 *in vitro*^49^, this has been implicated in a reduction in *P. aeruginosa* biofilm formation^50^. Similarly, the concentration of viable *P. aeruginosa* for the monomicrobial conditions did not show a reduction with the addition of 16HBE cells, indicating that the presence of *S. aureus* may have impacted hBDs expression.

When 16HBE cells were included within the airway models, monomicrobial *P. aeruginosa* tended to show a greater resistance to ciprofloxacin when grown within the healthy airway model, although the average concentration of viable *P. aeruginosa* within the hydrogels was very similar for the healthy and CF models (Figure 4B). Along with this trend, the average growth of *P. aeruginosa* within the healthy model appeared to increase with the addition of 16HBE cells while the average growth of *P. aeruginosa* within the CF model was similar with and without 16HBE cells. Other airway model studies have also reported changes in *P. aeruginosa* behavior when grown on lung epithelial cells, such as altered gene expression or an increase in antibiotic tolerance^35,51^. A potential cause of increased *P. aeruginosa* growth in the healthy model could be due to the influence of epithelial cell metabolites^52,53^. While host-microbe metabolic interactions in CF airway infections have yet to be fully understood due to the complex and polymicrobial nature of these infections, there is evidence that the effectiveness of antibiotics can be modulated by host metabolites^54^ such as lactate^55^.

When 16HBE cells were included within the polymicrobial models, *S. aureus* tended to show a greater resistance to ciprofloxacin within the CF airway model as compared to the healthy airway model. This trend was more consistent than what was observed when 16HBE cells were not included (Figure 4G and 4H). Co-culture with *P. aeruginosa* has been shown to increase *S. aureus* surface protein expression, leading to greater surface binding^56^. The increased resistance of *S. aureus* to ciprofloxacin within the polymicrobial CF airway model could also be attributed to the possible formation of dual-species biofilms with *P. aeruginosa*, which has been found in previous studies^45,57^. Overall, few trends were observed for the growth of *S. aureus* in the mono- and poly-microbial growth conditions other than what is discussed above. One possible explanation for the lack of trends in *S. aureus* culture was the use of a potentially subinhibitory ciprofloxacin concentration for the species. While 0.5 μg/ml ciprofloxacin reduces *S. aureus* growth relative to without ciprofloxacin, this concentration appears to potentially be subinhibitory within the airway models despite being greater than the MIC when tested in conditions without the mucus-like hydrogels. This subinhibitory concentration could have induced changes in *S. aureus*, such as increased virulence and toxin secretion, which has been found to occur with *S. aureus* and other antibiotics^58^.

The viability of 16HBE cells was assessed for multiple culture conditions, including within the healthy and CF airway models (Figure 1) with and without ciprofloxacin. Although 16HBE cells were non-viable for all of the culture conditions at the 24-hour timepoint with no ciprofloxacin, cell morphology and staining differed between *P. aeruginosa* and *S. aureus* mono- and polymicrobial conditions at this timepoint. For the monomicrobial *P. aeruginosa* condition with no ciprofloxacin, all 16HBE cells had detached from the surface of the wells. Other *in vitro* studies have found that *P. aeruginosa* causes epithelial cell detachment in as short as 3 hours of co-culture^19,35,51,59^. The type II and type III secretion systems have both been implicated in *P. aeruginosa* virulence and host cell cytotoxicity^35,60,61^ and 16HBE cell detachment observed in the 24-hour monomicrobial *P. aeruginosa* was likely due to toxins secreted by these systems.

When the 16HBE cells were grown within the airway models in the presence of 0.5 μg/ml ciprofloxacin, their viability was assessed after 48 hours of culture (Figure 5). For the monomicrobial *P. aeruginosa*, 16HBE cell viability was much more consistent and appeared to be relatively high for all three biological replicates, although 16HBE cell attachment was generally reduced within the CF airway model compared to the healthy model. This cell detachment indicates that there was likely some 16HBE killing by *P. aeruginosa*^19,35,51,59^ but to a lesser extent than what was observed in the airway models with no ciprofloxacin. The inclusion of 0.5 μg/ml ciprofloxacin within the airway models provided some control of bacteria growth which resulted in variable cell viability depending on bacteria species.

When 16HBE cells were grown with monomicrobial *S. aureus* without ciprofloxacin, all 16HBE cells were found to be non-viable at the 24-hour time point yet exhibited differing morphology as compared to the 16HBE cells killed in the monomicrobial *P. aeruginosa* condition. For monomicrobial *S. aureus* conditions, the 16HBE cells remained adherent but were non-viable. This 16HBE cell killing could be due to potential epithelial cell internalization and subsequent intracellular replication of *S. aureus*, which have been found to cause host cell cytotoxicity including the induction of apoptosis^62–64^.

The CF airway model demonstrated multiple features characteristic of chronic CF airway infections. One of the features of chronic CF airway infections is an increased tolerance to antibiotics. The ciprofloxacin MIC was determined to be 0.125 μg/ml for monomicrobial *P. aeruginosa* and *S. aureus* without the presence of a mucus-like hydrogel or mammalian cells. The mono- and polymicrobial *P. aeruginosa* and *S. aureus* remained viable for 48 hours within the CF airway model in the presence of 0.5 μg/ml ciprofloxacin, a concentration 4-fold greater than the MIC. An additional feature of chronic CF airway infections is altered or reduced virulence of *P. aeruginosa* which influences interactions between *P. aeruginosa* and *S. aureus*. *P. aeruginosa* strains isolated from long-term chronically infected patients have been shown to have a greater ability to co-exist with *S. aureus in vitro* relative to reference strains or early-adapted strains^65,66^. Many models lack the ability to capture the coexistence of *P. aeruginosa* and *S. aureus in vitro* despite it being known that the two co-infect CF patient airways *in vivo*^14,67^. In the CF airway model described here we were able to sustain *P. aeruginosa* and *S. aureus* in polymicrobial growth conditions co-cultured with mammalian cells for 48 hours.

The longevity of the CF airway model is also a feature of its chronicity; this model, which remains stable for 48 hours, surpasses the culture periods achieved in other *in vitro* CF airway infection models which generally range from 2 to 24 hours^20,35–38^. With optimized ciprofloxacin concentrations, it is possible to reduce inter-replicate variation and extend the viability of the model beyond 48 hours. Nonetheless, the increased culture period achieved for the CF airway model in this study is beneficial to its use for studying longer term growth behavior of CF pathogens.

## Methods

### Hydrogel preparation

Stock sodium alginate solutions were prepared at a concentration of 30 mg/ml alginic acid sodium salt (low viscosity, MP Biomedicals) in PBS. Alginic acid solution was sterilized via UV sterilization. HEPES buffer was added to the stock sodium alginate solution at a concentration of 75 mM. Stock CaCl_2_ (10mg/ml) solution was diluted with 0.9% NaCl in DI water and sterile filtered prior to use. Sterilized Mucin Type II (PGM, Sigma-Aldrich) powder was solubilized with PBS to create a 100 mg/mL solution. To prepare the hydrogels, alginate and mucin solutions were first mixed and diluted with PBS. The CaCl_2_ stock solutions were then diluted with the NaCl solution and rapidly mixed with the rest of the hydrogel components. Hydrogel formulations tested were combinations of 1% and 1.5% (w/v) alginate, 0.5 mg/ml and 0.6 mg/ml CaCl_2_ crosslinker, and 0%, 1%, and 4% (w/v) mucins. The hydrogel mixture was then deposited into the wells at a thickness of 1.5 mm (based on volume deposited) and allowed to solidify within a cell culture incubator at 37°C and 5% CO_2_ for 45 minutes. A thin layer of CaCl_2_-crosslinked alginate hydrogel (alginate cushion) with 0% mucin concentration was also incorporated into the model. The alginate cushion was prepared with the same steps as outlined above with 1.5% alginate and 0.5 mg/ml CaCl_2_ crosslinker.

### Mammalian cell culture

The 16HBE cell line (16HBE14o-) was kindly provided by Dr. Geoff Maksym, Dalhousie University. 16HBE cells were incubated at 37°C and 5% CO_2_. Dulbecco’s Modified Eagle Medium (DMEM, Thermo Fisher Scientific, Gibco, Cat. No. 11965092) and Gibco Ham’s F-12 nutrient mixture (F-12, Thermo Fisher Scientific, Gibco, Cat. No. 11765054) were mixed at a 1:1 ratio and supplemented with 10% fetal bovine serum (FBS, Thermo Fisher Scientific, Gibco) and 1% antibiotic-antimycotic (AA, 100x, Thermo Fisher Scientific, Gibco, Cat. No. 15240062) for all 16HBE cell growth and maintenance. 16HBE cells were seeded at a cell density of approximately 50000 to 55000 cells/well in a 48-cell cell culture plate. Wells were coated with fibronectin (Thermo Fisher Scientific) prior to cell seeding to encourage cell adhesion. The cells were grown for 24 hours in the 48-well plate prior to use for experiments.

### Cytotoxicity assessment of hydrogels

To assess the cytotoxicity of potential hydrogel formulations, 16HBE cells were first cultured as stated above. The cushion hydrogel was first added to the individual wells at a thickness of approximately 0.4 mm and allowed to crosslink within a cell culture incubator at 37°C and 5% CO_2_ for 10 minutes. The mucus-like hydrogel formulations were then added over top of the cushion at a thickness of approximately 1.1 mm and allowed to solidify in the incubator for 35 minutes. After the hydrogels were solidified, 250 µl of cell culture medium was added over top of the hydrogels. The cells and hydrogels were then cultured together for 48 hours at 37°C and 5% CO_2_. After 48 hours, the cell culture medium and hydrogels were removed.

Live/dead assay consisting of calcein AM and ethidium homodimer-1 (Thermo Fisher Scientific) and a Hoechst stain (Hoechst 33342, Thermo Fisher Scientific) was performed to assess cell viability as per manufacturers protocols. EVOS™ FL Auto 2 Imaging System (Thermo Fisher Scientific) was used to collect phase contrast and fluorescence images and to visualize live cells (green fluorescent protein channel, GFP), dead cells (red fluorescent protein channel, RFP), and cellular DNA (DAPI channel). ImageJ was used for image processing and cell viability quantification. Cell viability for each condition was calculated based on cell counts using Equation 1.

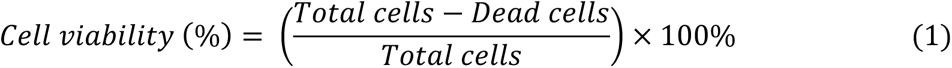

### Viscoelastic characterization of hydrogels

The storage (elastic, G’) and loss (viscous, G’’) moduli of the mucus-like hydrogel formulations were measured. A controlled-stress/controlled-rate rheometer (AR2000, TA Instruments) with a 40 mm diameter/2° cone geometry was used to measure the moduli. Use of the rheometer was kindly provided by Dr. Kevin Plucknett and Dr. Gianfranco Mazzanti, Dalhousie University. Temperature was kept constant at 23°C using a Peltier plate and a solvent trap was used to prevent changes in hydrogel moisture content. Strain sweeps were performed to determine the linear viscoelastic region (LVR) of each hydrogel. Strain sweeps were performed from 0.1% to 1000% strain at an angular frequency of 1 rad/s. Frequency sweeps were then performed to determine the frequency-dependent storage modulus and loss modulus of each hydrogel. Frequency sweeps were performed from 0.1 to 100 rad/s at strains found to be within the LVR of each hydrogel, which ranged from 1% to 2% strain. Each hydrogel formulation was tested three times. To conduct the measurements, the hydrogels were first prepared as outlined above and 590 µl was deposited onto the Peltier plate. Hydrogels were deposited directly after mixing the crosslinker and tests were conducted 15 or 20 minutes after deposition to allow the hydrogels to solidify. Strain and frequency sweeps were repeated three times for each hydrogel formulation.

### Bacteria strains and culture conditions

*P. aeruginosa* CF18 and *S. aureus* ATCC 6538 were used for this study. *P. aeruginosa* CF18 is a CF clinical strain isolated from the airway of a child (< 24 months)^68^ with CF and was kindly provided by Dr. Zhenyu Cheng, Dalhousie University. Frozen stocks of *P. aeruginosa* CF18 were stored in Luria-Bertani (LB) broth containing 25% (v/v) glycerol stored at -80°C. For use of the bacteria within experiments, frozen stocks were streaked on 1.5% (w/v) LB agar (Sigma-Aldrich) plates and incubated overnight at 37°C, 5% CO_2_ for 16 to 18 hours. LB broth was used for *P. aeruginosa* overnight cultures harvested from streaked plates and Brain Heart Infusion (BHI) broth was used for *S. aureus* overnight cultures. The inoculated broth was placed in a shaking incubator shaking at 200 rpm and 37°C for 16 to 18 hours.

### Preparation of ATPS

The ATPS formulations used for this research consisted of polyethylene glycol (PEG) and dextran (DEX). To prepare the ATPS formulations, 5% (w/v) PEG (35 kDa, Sigma-Aldrich) and 5% (w/v) DEX (500 kDa, Pharmacosmos) were dissolved together in LB broth as well as in DMEM/F-12 cell culture medium supplemented with 10% FBS. The cell culture medium used for ATPS preparation did not contain any antibiotics or antimycotics. The solutions were then mixed on a rocking platform shaker (VWR) until the PEG and DEX were fully dissolved. After sterile filtration, the solutions were centrifuged for 90 minutes at 3000 x g to separate the PEG-rich phase and the DEX-rich phase, which were then collected and stored at 4°C. The ATPS used was a combination of the PEG-rich phase of the DMEM/F-12 ATPS formulation (to support 16HBE cell growth) with the DEX-rich phase from the LB broth ATPS formulation (to support microbe growth).

### Assessment of short-term bacteria culture establishment over healthy and CF hydrogels using an aqueous two-phase system

A short-term confined bacteria culture was established over top of the healthy and CF hydrogels using the ATPS. The alginate cushion and the healthy and CF mucus-like hydrogels were first deposited into wells of a 48-well plate at a total height of approximately 1.5 mm (based on volume deposited). 250 µl of the PEG-rich phase from a 5%/5% PEG/DEX ATPS in DMEM/F-12 was then added over top of the hydrogels. *P. aeruginosa* CF18 and *S. aureus* ATCC 6538 from overnight cultures were suspended in monoculture and co-culture in the DEX-rich phase of a 5%/5% PEG/DEX ATPS in cell culture media. The bacteria monocultures were suspended within the DEX-rich phase at an OD_600_ of 0.01 and the bacteria co-cultures were suspended within the DEX-rich phase at an OD_600_ of 0.005 for each species giving a total OD_600_ of 0.01. Establishment of the ATPS was accomplished as previously described^69^. The plate was then incubated for 5 hours at 37°C and 5% CO_2_. After 5 hours, the liquid PEG phase and the hydrogel phase of each culture condition were removed separately. Both phases were vortexed for 60 seconds and centrifuged at 16000 x g for 10 minutes to pellet bacteria. After centrifugation, the supernatant from each condition was removed, the hydrogel phases were resuspended in 150 µl of PBS, and the PEG phases were resuspended in 250 µl. The samples were then serially diluted with PBS. Samples from monoculture conditions were diluted from 10^-1^ to 10^-6^ and 20 µl of each dilution was spot plated onto LB agar (for *P. aeruginosa* monoculture) and BHI agar (for *S. aureus* monoculture). The samples from co-culture conditions were then diluted from 10^-2^ to 10^-5^ for PEG phase samples and 10^-4^ to 10^-7^ for hydrogel samples and 100 µl of each dilution was spread plated onto LB agar. *P. aeruginosa* CF18 and *S. aureus* ATCC 6538 were distinguishable based on colony color and morphology. Agar plates were incubated for 16 to 18 hours at 37°C, and 5% CO_2_ and colonies were counted for colony forming unit (CFU) determination within the PEG phase and hydrogel phase of each condition. Equation 2 was used to calculate the concentration of viable bacteria within each condition.

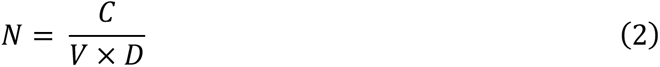

In Equation 2, N is the concentration of viable bacteria in CFU/ml, C is the number of colonies counted, V is the volume of sample plated in ml, and D is the dilution factor.

### Determination of ciprofloxacin minimum inhibitory concentration for bacteria monocultures and co-cultures

The minimum inhibitory concentration (MIC) of ciprofloxacin was determined for *P. aeruginosa* CF18 and *S. aureus* ATCC 6538 in monoculture and in co-culture using a broth microdilution procedure as previously described^70^. Prior to this procedure, the correlation between OD_600_ and CFU/ml was first determined for *P. aeruginosa* CF18 and *S. aureus* ATCC 6538. Overnight cultures of *P. aeruginosa* and *S. aureus* were diluted to an OD_600_ of 0.01, serially diluted in PBS from 10^-4^ to 10^-7^, and 100 µl of each dilution was spread plated on LB agar. The plates were incubated at 37°C and 5% CO_2_ for 16 to 18 hours. After incubation, the colonies were counted and the CFU/ml corresponding to the plated OD_600_ of 0.01 was calculated using Equation 2. This was repeated in triplicate using overnight cultures from three separate colonies for both species of bacteria.

### Assessment of healthy and CF airway models

Prior to culturing bacteria within the airway models, the growth of the bacteria with the mucus-like hydrogels in the presence of ciprofloxacin was first assessed with and without the inclusion of 16HBE cells. The cultures were prepared and deposited on the gels via ATPS as described above. The plate was cultured for 5 hours at 37°C and 5% CO_2_. After 5 hours, the PEG-rich phase of the ATPS was removed through pipetting and was replaced with cell culture media supplemented with 0.5 µg/ml ciprofloxacin. The plate was then cultured for an additional 43 hours at 37°C and 5% CO_2_, for a total culture period of 48 hours. Then, 50 µl of the liquid phase from each culture condition was collected and the remainder of the liquid phase was disposed of. This was done to avoid collecting any of the bacteria at the liquid-hydrogel interface and to avoid collecting the hydrogel itself. Once the liquid phase was removed, the hydrogel phase from each culture condition was collected and CFU was determined as described above. Samples from co-culture conditions were diluted from 10^-1^ to 10^-6^ and 100 µl of each dilution was spread plated onto LB agar. For culture conditions with 16HBE, after the 48-hour culture period, cell viability was assessed using the live/dead assay and a Hoechst stain as described in above. A schematic of the final assessment of the airway models can be found in Figure S3.

### Statistical analysis

GraphPad Prism (Version 9.4.1) was used for statistical analysis of bacterial abundance and potential mucus-like hydrogel rheology data. Statistical significance of bacterial abundance was determined using two-way analysis of variance (ANOVA) with p values represented as *p < 0.05, **p < 0.01, ***p < 0.001. Statistical significance of potential mucus-like hydrogel rheology data when comparing increased crosslinker concentration and increased alginate concentration were determined using Welch’s t-test with p values represented as ^a^p < 0.05 (when comparing G’ values) and ^b^p < 0.05 (when comparing G’’ values). Statistical significance of potential mucus-like hydrogel rheology data when comparing increased mucin concentration was determined using one-way ANOVA with p values of p < 0.05 indicated by matching superscripts.

## Author Contributions

BL conceived the project. BL, COB, AH, AS and NJ designed and acquired the experiments. The manuscript was written through contributions of BL, COB, SS, NJ, AS, AH, and ZC. All authors have given approval to the final version of the manuscript.

## References

1. Wagner, C. E., Wheeler, K. M. & Ribbeck, K. Mucins and their role in shaping the functions of mucus barriers. Annu. Rev. Cell Dev. Biol. 34, 189–215 (2018).

2. Fahy, J. V. & Dickey, B. F. Airway mucus function and dysfunction. N. Engl. J. Med. 363, 2233–2247 (2010).

3. Hill, D. B. et al. A biophysical basis for mucus solids concentration as a candidate biomarker for airways disease. PLoS One 9, e87681 (2014).

4. Henderson, A. G. et al. Cystic fibrosis airway secretions exhibit mucin hyperconcentration and increased osmotic pressure. J. Clin. Invest. 124, 3047–3060 (2014).

5. Burgel, P. R., Montani, D., Danel, C., Dusser, D. J. & Nadel, J. A. A morphometric study of mucins and small airway plugging in cystic fibrosis. Thorax 62, 153–161 (2007).

6. Ghanem, R. et al. Apparent yield stress of sputum as a relevant biomarker in cystic fibrosis. Cells 10, 3107 (2021).

7. Völler, M. et al. An optimized protocol for assessment of sputum macrorheology in health and muco-obstructive lung disease. Front. Physiol. 13, 912049 (2022).

8. Matsui, H. et al. Evidence for periciliary liquid layer depletion, not abnormal ion composition, in the pathogenesis of cystic fibrosis airways disease. Cell 95, 1005–1015 (1998).

9. Button, B. et al. A periciliary brush promotes the lung health by separating the mucus layer from airway epithelia. Science (80-.). 337, 937–941 (2012).

10. Button, B. et al. Roles of mucus adhesion and cohesion in cough clearance. Proc. Natl. Acad. Sci. U. S. A. 115, 12501–12506 (2018).

11. Kidd, T. J. et al. Defining antimicrobial resistance in cystic fibrosis. J. Cyst. Fibros. 17, 696–704 (2018).

12. Chmiel, J. F. et al. Antibiotic management of lung infections in cystic fibrosis: I. The microbiome, methicillin-resistant *Staphylococcus aureus*, gram-negative bacteria, and multiple infections. Ann. Am. Thorac. Soc. 11, 1120–1129 (2014).

13. Beck, J. M., Young, V. B. & Huffnagle, G. B. The microbiome of the lung. Transl. Res. 160, 258–266 (2012).

14. Millette, G. et al. Despite antagonism in vitro, *Pseudomonas aeruginosa* enhances *Staphylococcus aureus* colonization in a murine lung infection model. Front. Microbiol. 10, 2880 (2019).

15. Hoffmann, N. et al. Novel mouse model of chronic *Pseudomonas aeruginosa* lung infection mimicking cystic fibrosis. Infect. Immun. 73, 250–2514 (2005).

16. Pezzulo, A. A. et al. Reduced airway surface pH impairs bacterial killing in the porcine cystic fibrosis lung. Nature 487, 109–113 (2012).

17. Stoltz, D. A. et al. Cystic fibrosis pigs develop lung disease and exhibit defective bacterial eradication at birth. Sci. Transl. Med. 2, 29ra31 (2010).

18. Filkins, L. M. et al. Coculture of *Staphylococcus aureus* with *Pseudomonas aeruginosa* drives *S. aureus* towards fermentative metabolism and reduced viability in a cystic fibrosis model. J. Bacteriol. 197, 2252–2264 (2015).

19. Carterson, A. J. et al. A549 lung epithelial cells grown as three-dimensional aggregates: Alternative tissue culture model for *Pseudomonas aeruginosa* pathogenesis. Infect. Immun. 73, 1129–1140 (2005).

20. Orazi, G. & O’Toole, G. A. *Pseudomonas aeruginosa* alters *Staphylococcus aureus* sensitivity to vancomycin in a biofilm model of cystic fibrosis infection. MBio 8, e00873–17 (2017).

21. Limoli, D. H., et al. *Pseudomonas aeruginosa* alginate overproduction promotes coexistance with *Staphylococcus aureus* in a model of cystic fibrosis respiratory infection. Am. Soc. Microbiol. 8, e00186–17 (2017).

22. Sriramulu, D. D., Lünsdorf, H., Lam, J. S. & Römling, U. Microcolony formation: A novel biofilm model of *Pseudomonas aeruginosa* for the cystic fibrosis lung. J. Med. Microbiol. 54, 667–676 (2005).

23. Frisch, S. et al. A pulmonary mucus surrogate for investigating antibiotic permeation and activity against *Pseudomonas aeruginosa* biofilms. J. Antimicrob. Chemother. 76, 1472– 1479 (2021).

24. Matsui, H. et al. A physical linkage between cystic fibrosis airway surface dehydration and *Pseudomonas aeruginosa* biofilms. Proc. Natl. Acad. Sci. U. S. A. 103, 18131–18136 (2006).

25. Deschamps, E. et al. Membrane phospholipid composition of *Pseudomonas aeruginosa* grown in a cystic fibrosis mucus-mimicking medium. Biochim. Biophys. Acta - Biomembr. 1863, 183482 (2021).

26. Haley, C. L., Colmer-Hamood, J. A. & Hamood, A. N. Characterization of biofilm-like structures formed by *Pseudomonas aeruginosa* in a synthetic mucus medium. BMC Microbiol. 12, 181 (2012).

27. Palmer, K. L., Aye, L. M. & Whiteley, M. Nutritional cues control *Pseudomonas aeruginosa* multicellular behavior in cystic fibrosis sputum. J. Bacteriol. 189, 8079–8087 (2007).

28. Huck, B. C. et al. Macro- and microrheological properties of mucus surrogates in comparison to native intestinal and pulmonary mucus. Biomacromolecules 20, 3504–3512 (2019).

29. Darch, S. E. et al. Spatial determinants of quorum signaling in a *Pseudomonas aeruginosa* infection model. Proc. Natl. Acad. Sci. U. S. A. 115, 4779–4784 (2018).

30. Hamed, R. & Fiegel, J. Synthetic tracheal mucus with native rheological and surface tension properties. Soc. Biomater. 102A, 1788–1798 (2013).

31. Hansen, I. M. et al. Hyaluronic acid molecular weight-dependent modulation of mucin nanostructure for potential mucosal therapeutic applications. Mol. Pharm. 14, 2359–2367 (2017).

32. Pacheco, D. P. et al. Disassembling the complexity of mucus barriers to develop a fast screening tool for early drug discovery. J. Mater. Chem. B 7, 4940–4952 (2019).

33. Butnarasu, C., Caron, G., Pacheco, D. P., Petrini, P. & Visentin, S. Cystic fibrosis mucus model to design more efficient drug therapies. Mol. Pharm. 19, 520–531 (2022).

34. Huang, A. J. et al. Characterization of an engineered mucus microenvironment for in vitro modeling of host–microbe interactions. Sci. Rep. 12, 5515 (2022).

35. Anderson, G. G., Moreau-Marquis, S., Stanton, B. A. & O’Toole, G. A. In vitro analysis of tobramycin-treated *Pseudomonas aeruginosa* biofilms on cystic fibrosis-derived airway epithelial cells. Infect. Immun. 76, 1423–1433 (2008).

36. Briaud, P. et al. Coexistence with *Pseudomonas aeruginosa* alters *Staphylococcus aureus* transcriptome, antibiotic resistance and internalization into epithelial cells. Sci. Rep. 9, 16564 (2019).

37. Crabbé, A. et al. Antimicrobial efficacy against *Pseudomonas aeruginosa* biofilm formation in a three-dimensional lung epithelial model and the influence of fetal bovine serum. Sci. Rep. 7, 43321 (2017).

38. Yu, Q. et al. In vitro evaluation of tobramycin and aztreonam versus *Pseudomonas aeruginosa* biofilms on cystic fibrosis-derived human airway epithelial cells. J. Antimicrob. Chemother. 67, 2673–2681 (2012).

39. Boon, M. et al. Morphometric analysis of explant lungs in cystic fibrosis. Am. J. Respir. Crit. Care Med. 193, 516–526 (2016).

40. Nielsen, H., Hvidt, S., Sheils, C. A. & Janmey, P. A. Elastic contributions dominate the viscoelastic properties of sputum from cystic fibrosis patients. Biophys. Chem. 112, 193– 200 (2004).

41. Dawson, M., Wirtz, D. & Hanes, J. Enhanced viscoelasticity of human cystic fibrotic sputum correlates with increasing microheterogeneity in particle transport. J. Biol. Chem. 278, 50393–50401 (2003).

42. Schuster, B. S., Soo, J., Woodworth, G. F. & Hanes, J. Nanoparticle diffusion in respiratory mucus from humans without lung disease. Biomaterials 34, 3439–3446 (2013).

43. Markovetz, M. R. et al. Endotracheal tube mucus as a source of airway mucus for rheological study. Am. J. Physiol. Lung Cell. Mol. Physiol. 317, L498–L509 (2019).

44. Beaudoin, T., et al. *Staphylococcus aureus* interaction with *Pseudomonas aeruginosa* biofilm enhances tobramycin resistance. npj Biofilms Microbiomes 3, 25 (2017).

45. Alves, P. M. et al. Interaction between *Staphylococcus aureus* and *Pseudomonas aeruginosa* is beneficial for colonisation and pathogenicity in a mixed biofilm. Pathog. Dis. 76, fty003 (2018).

46. Alhede, M. et al. Phenotypes of non-attached *Pseudomonas aeruginosa* aggregates resemble surface attached biofilm. PLoS One 6, e27943 (2011).

47. Pazgier, M., Hoover, D. M., Yang, D., Lu, W. & Lubkowski, J. Human β-defensins. Cell. Mol. Life Sci. 63, 1294–1313 (2006).

48. Alekseeva, L. et al. Inducible expression of beta defensins by human respiratory epithelial cells exposed to *Aspergillus fumigatus* organisms. BMC Microbiol. 9, 33 (2009).

49. Midorikawa, K., et al. *Staphylococcus aureus* susceptibility to innate antimicrobial peptides, β-defensins and CAP18, expressed by human keratinocytes. Infect. Immun. 71, 3730–3739 (2003).

50. Parducho, K. R. et al. The antimicrobial peptide human beta-defensin 2 inhibits biofilm production of *Pseudomonas aeruginosa* without compromising metabolic activity. Front. Immunol. 11, (2020).

51. Moreau-Marquis, S. et al. The ΔF508-CFTR mutation results in increased biofilm formation by *Pseudomonas aeruginosa* by increasing iron availability. Am. J. Physiol. - Lung Cell. Mol. Physiol. 295, 25–37 (2008).

52. Crabbé, A., Jensen, P. Ø., Bjarnsholt, T. & Coenye, T. Antimicrobial tolerance and metabolic adaptations in microbial biofilms. Trends Microbiol. 27, 850–863 (2019).

53. Jean-Pierre, F., Henson, M. A. & O’Toole, G. A. Metabolic modeling to interrogate microbial disease: A tale for experimentalists. Front. Mol. Biosci. 8, 634479 (2021).

54. Crabbé, A. et al. Host metabolites stimulate the bacterial proton motive force to enhance the activity of aminoglycoside antibiotics. PLoS Pathog. 15, e1007697 (2019).

55. Wang, Y. et al. Two NAD-independent l-lactate dehydrogenases drive l-lactate utilization in *Pseudomonas aeruginosa* PAO1. Environ. Microbiol. Rep. 10, 569–575 (2018).

56. Kumar, A. & Ting, Y. P. Presence of *Pseudomonas aeruginosa* influences biofilm formation and surface protein expression of *Staphylococcus aureus*. Environ. Microbiol. 17, 4459–4468 (2015).

57. Magalhães, A. P., Lopes, S. P. & Pereira, M. O. Insights into cystic fibrosis polymicrobial consortia: The role of species interactions in biofilm development, phenotype, and response to in-use antibiotics. Front. Microbiol. 7, 2146 (2017).

58. Gao, P. et al. Subinhibitory concentrations of antibiotics exacerbate staphylococcal infection by inducing bacterial virulence. Microbiol. Spectr. 10, e0064022 (2022).

59. Chi, E., Mehl, T., Nunn, D. & Lory, S. Interaction of *Pseudomonas aeruginosa* with A549 pneumocyte cells. Infect. Immun. 59, 822–828 (1991).

60. Jyot, J. et al. Type II secretion system of *Pseudomonas aeruginosa*: In vivo evidence of a significant role in death due to lung infection. J. Infect. Dis. 203, 1369–1377 (2011).

61. Horna, G. & Ruiz, J. Type 3 secretion system of *Pseudomonas aeruginosa*. Microbiol. Res. 246, 126719 (2021).

62. Korea, C. G. et al. Staphylococcal Esx proteins modulate apoptosis and release of intracellular *Staphylococcus aureus* during infection in epithelial cells. Infect. Immun. 82, 4144–4153 (2014).

63. Stelzner, K. et al. Intracellular *Staphylococcus aureus* employs the cysteine protease staphopain A to induce host cell death in epithelial cells. PLoS Pathog. 17, e1009874 (2021).

64. Kahl, B. C. et al. *Staphylococcus aureus* RN6390 replicates and induces apoptosis in a pulmonary epithelial cell line. Infect. Immun. 68, 5385–5392 (2000).

65. Michelsen, C. F., et al. *Staphylococcus aureus* alters growth activity, autolysis, and antibiotic tolerance in a human host-adapted *Pseudomonas aeruginosa* lineage. J. Bacteriol. 196, 3903–3911 (2014).

66. Baldan, R. et al. Adaptation of *Pseudomonas aeruginosa* in cystic fibrosis airways influences virulence of *Staphylococcus aureus* in vitro and murine models of co-infection. PLoS One 9, e89614 (2014).

67. Cystic Fibrosis Foundation. 2019 Patient Registry Annual Data Report. (2020).

68. Wolfgang, M. C. et al. Conservation of genome content and virulence determinants among clinical and environmental isolates of *Pseudomonas aeruginosa*. Proc. Natl. Acad. Sci. U. S. A. 100, 8484–8489 (2003).

69. Huang, A. J., Clarke, A. N., Jafari, N. & Leung, B. M. Characterization of patterned microbial growth dynamics in aqueous two-phase polymer scaffolds. ACS Biomater. Sci. Eng. 7, 5506–5514 (2021).

70. Wiegand, I., Hilpert, K. & Hancock, R. E. W. Agar and broth dilution methods to determine the minimal inhibitory concentration (MIC) of antimicrobial substances. Nat. Protoc. 3, 163–175 (2008).

