## Supplemental Data for "Modeling Cystic Fibrosis Chronic Infection Using Engineered Mucus-like Hydrogels"

| Gel name | [Mucin] (%) | [Alginate] (%) | $[CaCl_2]$ (mg/ml) | G' (Pa) | G'' (Pa) |
|---|---|---|---|---|---|
| A | 0 | 1 | 0.5 | 13.600 ± 3.170 | 1.300 ± 0.382 |
| B | 1 | 1 | 0.5 | 10.039 ± 3.127 | 1.326 ± 0.351 |
| C | 4 | 1 | 0.5 | 10.729 ± 2.907 | 2.759 ± 0.555 |
| D | 0 | 1 | 0.6 | 15.596 ± 9.801 | 2.030 ± 0.554 |
| E | 1 | 1 | 0.6 | 11.118 ± 3.571 | 2.902 ± 0.761 |
| F | 4 | 1 | 0.6 | 16.590 ± 5.801 | 4.998 ± 1.733 |
| G | 0 | 1.5 | 0.5 | 30.283 ± 9.229 | 2.400 ± 0.381 |
| H | 1 | 1.5 | 0.5 | 22.830 ± 4.826 | 3.003 ± 1.010 |
| I | 4 | 1.5 | 0.5 | 17.463 ± 3.535 | 3.654 ± 0.797 |
| J | 0 | 1.5 | 0.6 | 34.593 ± 8.580 | 3.224 ± 0.413 |
| K | 1 | 1.5 | 0.6 | 19.970 ± 4.826 | 3.224 ± 0.798 |
| L | 4 | 1.5 | 0.6 | 19.970 ± 2.523 | 5.817 ± 0.260 |

**Table S1:** *Storage (G') and loss (G'') moduli of potential mucus-like hydrogel formulations at an angular frequency of 1 rad/s presented in this table.* Values are the average ± SD from three independent measurements. Gel-E was chosen as the healthy mucus-like hydrogel formulation and Gel-L was chosen as the CF mucus-like hydrogel formulation. The percent solids (i.e. the concentration of mucins and alginate) as well as the crosslinker concentration were found to impact the viscoelastic properties of the hydrogels.

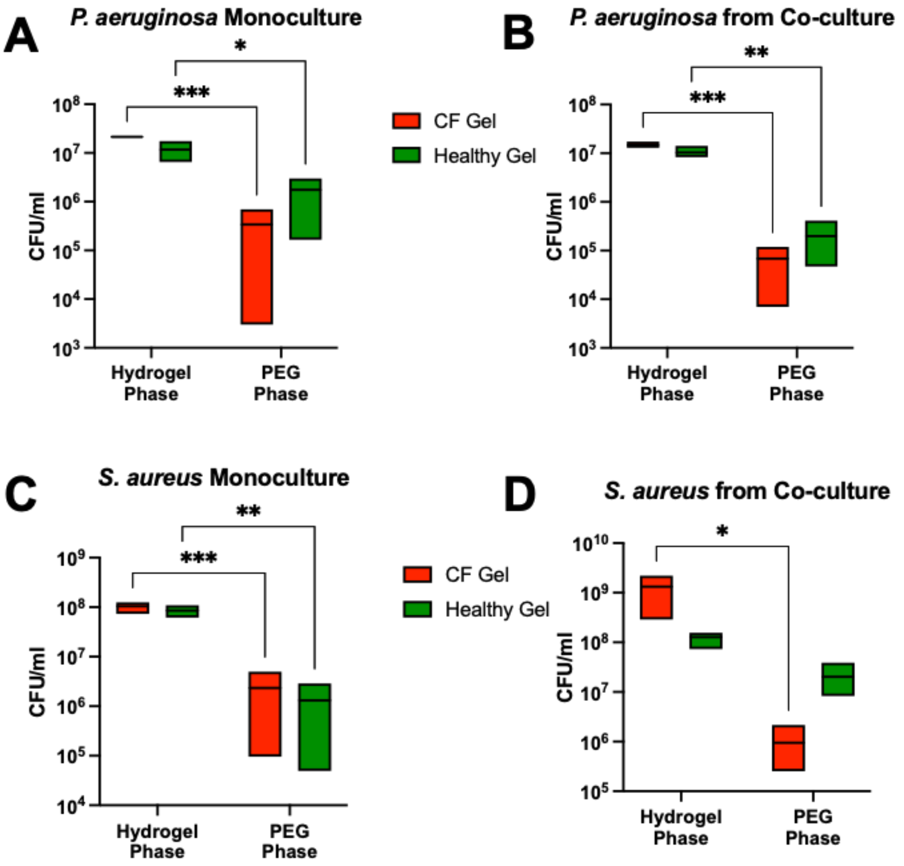


**Figure S1: Concentration of viable bacteria (CFU/ml) within the PEG phase of the ATPS (i.e. the less dense liquid phase) and the hydrogel phase after short-term (5 hour) monoculture and co-culture to demonstrate the ability to establish a confined culture over the hydrogels.** *P. aeruginosa* grown in (A) monoculture and *P. aeruginosa* grown in (B) co-culture with *S. aureus*. *S. aureus* grown in (C) monoculture and *S. aureus* grown in (D) co-culture with *P. aeruginosa*. The bars span from the minimum to the maximum value with the line within the bars representing the average value from three biological replicates (n = 3). P values obtained using two-way ANOVA. *$p < 0.05$, **$p < 0.01$, ***$p < 0.001$.

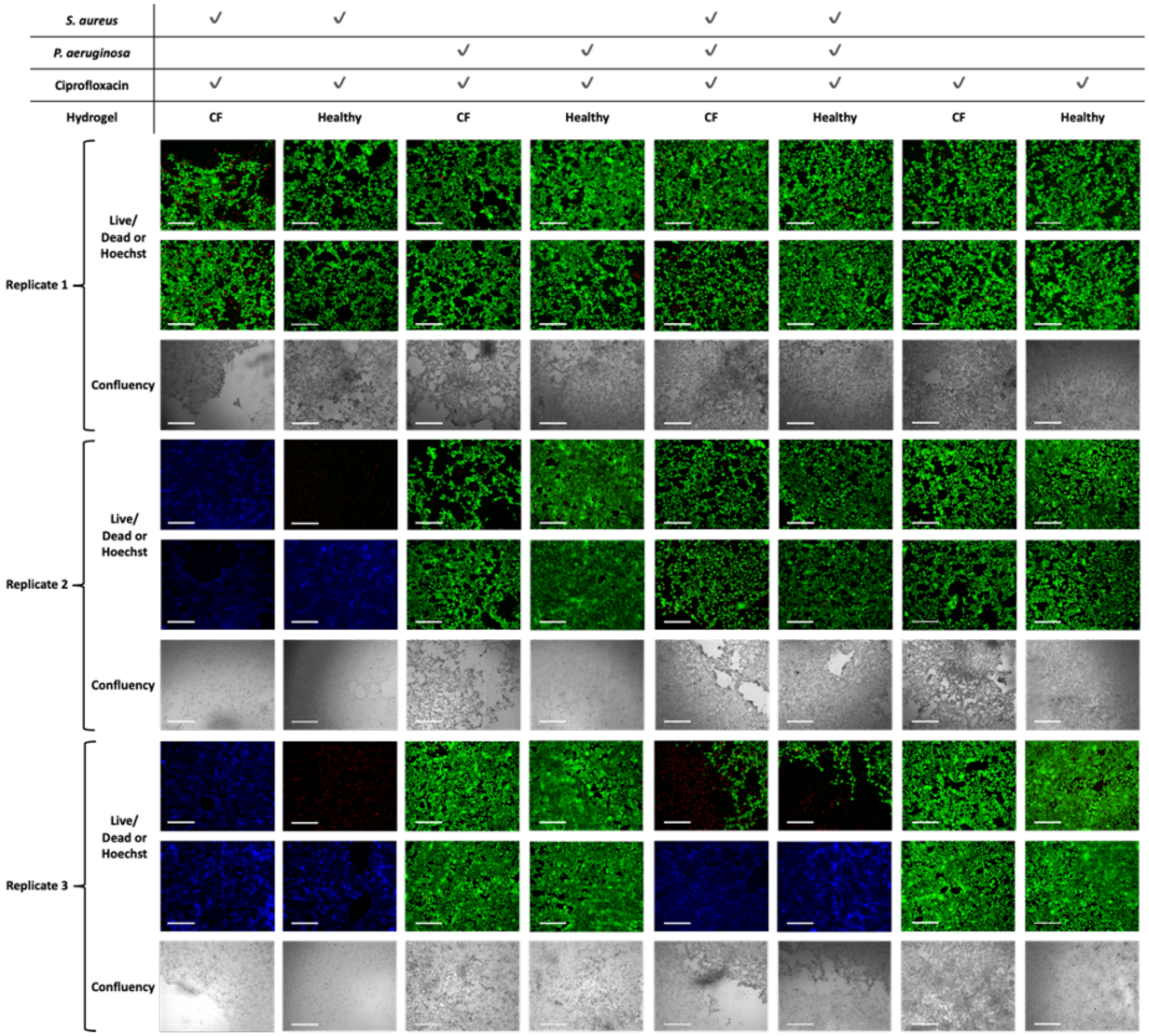


**Figure S2:** *16-HBE cell viability and overall adherence after culture within the CF and healthy airway models.* Conditions included *S. aureus* monoculture, *P. aeruginosa* monoculture, *P. aeruginosa* and *S. aureus* co-culture, and models with no bacteria. All conditions were exposed to 0.5 µg/ml ciprofloxacin. Live/dead assay was performed after 48 hours of culture. Green cells are viable and red cells are non-viable. Hoechst DNA stain (blue) presented in conditions where cellular material is remaining but is not staining green or red. Images from three biological replicates presented individually to capture variation between trials. Representative images selected from each replicate. Scale bar is 275 µm for fluorescent images (live/dead or Hoechst). Scale bar is 650 µm for cell confluency images.

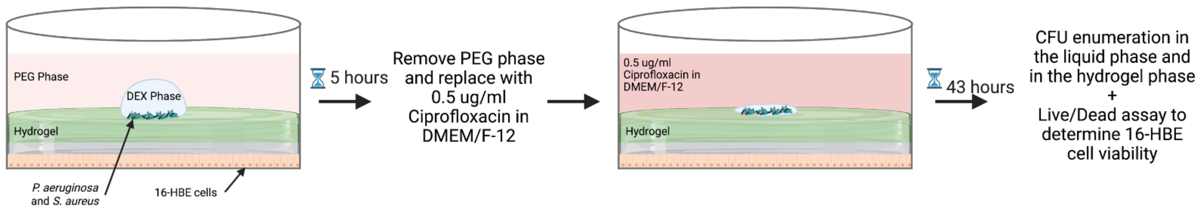


**Figure S3:** *Schematic of the final airway model assessment.* For the final assessment, both the healthy and CF airway models were composed of a mucus-like hydrogel overlaid onto a layer of 16HBE cells. An ATPS was used to deposit P. aeruginosa and S. aureus, monomicrobial or polymicrobial culture conditions, over top of the mucus-like hydrogels. The models were cultured for 5 hours and then the PEG phase of the ATPS was removed and replaced with 0.5 μg/ml ciprofloxacin in DMEM/F-12. This was then cultured for another 43 hours and, after this culture period, the concentrations of viable bacteria within the liquid and hydrogel phases were determined through CFU enumeration and 16HBE cell viability was assessed using a liv/dead assay and DNA stain.
